# The difficulties of predicting evolutionary response to selection

**DOI:** 10.1101/2023.02.01.526560

**Authors:** Mats Björklund

**Author notes:** Author contribution: MB did all the work. Data accessibility: No data to share.

## Abstract

There has been a long debate about the difficulties of predicting evolutionary change despite knowledge of selection and genetic variance. One reason might be the stochastic effects of sexual reproduction creating a variance of offspring genotypes and thus phenotypes. This was tested by means of an explicit genetic individual-based simulation with one trait determined by 50 loci. After 100 generations of weak fluctuating selection an experiment was performed where the optimum was displaced one standard deviation. The response and the predicted response were then compared for 100 populations. The simulation shows two things, first there is a considerable variation in response between populations. This was also found when the same population was replicated many times with the same selection. Second, using the individual-based estimates of genetic variance seriously overestimates the predicted response, while an approach using the variance of mean pair breeding values gave essentially unbiased predictions. However, in all cases there was a lack of precision. Hence, the “missing response” is, at least partly, due to an overestimation of predicted response and an inevitable variance around the predicted value.

---

Evolutionary biology is to a very large extent retrospective and descriptive trying to understand what has happened over evolutionary time and what kind of processes that has given rise to the patterns we now can observe. On the other hand, attempts to use current data to predict future evolution has been shown to be difficult. There are a number of studies where there is sufficient information about genetic variance and selection and thus a prediction should be possible. Despite this, these predictions have in many cases not been successful. In fact, of the examples listed in Walsh and Lynch (2018, Table 20.3) 22 out of 26 studies of natural populations failed to show a predicted response and there were even eight cases where the observed responses were in the opposite direction. Consequently, a great deal of effort has been devoted to understand this lack of match between observations and predictions (see e.g. Merilä et al. 2001a, Morrissey et al. 2012, Merilä and Hendry 2014, Pujol et al. 2018; Walsh and Lynch 2018). The factors discussed in these sources are obviously biologically relevant and there are good reasons to believe that they can account for some of the mismatch.

A factor that has not received much attention is the fact that in a sexually reproducing species the number of possible genotypes from a single mating pair is enormous. For example, as a result of recombination two parent of the same size can get offspring widely different in size, depending on the genotypes. The possible number of offspring genotypes increases very rapidly to astronomical numbers with the number of genes involved. This was discussed by Troy (2013) where he suggested that due to this factor alone, predicting evolution might be a very difficult task, in his words the evolution being an open-ended process with an infinite state space. This is might be an exaggeration but the vast number of possible genotypes that can result from a mating certainly needs to be considered in the discussion of the predicted response to selection.

In this paper I will test the impact of the capricious effect of recombination on the evolutionary response to a change in phenotypic optimum. I will use an individual-based simulation with a quantitative genetic trait determined by 50 loci. This approach allows for a comparison between estimated selection and response to the actual response. This will give important information on the predictability of evolutionary change, the accuracy and precision of the predicted response and the possibility of using different natural populations as replicates in experiments.

## Methods

I created a population with 500 diploid individuals each with 50 loci and two alleles with the values 1 and 2 randomly assigned to each allele. The allele frequencies were equal (50:50) at each locus across the population at start but due to this random assignment of allelic values individuals will differ in their genotypic setup. The genotypic value of each individual was then the sum of the allelic values, i.e. this mimics a polygenic trait with many loci with small effects. Phenotypes were created by adding a random variate from a normal distribution with a mean zero and a variance equal to the 1.5 times variance of the genotypes. This corresponds to the classic linear model where the phenotype of an individual is determined by the sum of the genotypic value and a random environmental value (e.g. Lynch and Walsh 1998). This creates a (broad-sense) heritability of around 0.5. Breeding values were calculated as in Lynch and Walsh (1998, p. 73). The variance of the breeding values is then the additive genetic variance. Since genotype frequencies are explicit in the calculation of breeding values, these were recalculated every generation since genotype frequencies change every generation.

The number of loci, individuals and number of generations were chosen such that mutations could not realistically play a role, which simplifies the analysis. Even if the mutation should be in the order of 10^−6^ per site per generation, the expected number of mutations would be five in total, ranging between zero and ten (number of mutations per generation = 1/(50 × 500 × 2) = 1/20, or one mutation per 20 generations). Hence, at any given time the frequency of a new mutation would be so small that it would not have any measurable effect. In addition, to have an effect the would mutation would have to be 1 → 0 or 2 → 3, while 1 → 2 and 2 → 1 would be basically neutral. If mutation rate is in the order of 10^−7^, which seems more realistic we could expect between zero and two mutations over a large set of replications, or on average one mutation every 200 generations.

At each generation fitness (W) of an individual was assigned by comparing the phenotype (z) to the optimum (θ),

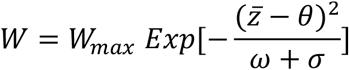

Where ω is the strength of stabilizing selection set to the phenotypic variance, and σ is the phenotypic variance, resulting in the total strength of stabilizing of around 80. This is a moderate strength of stabilizing selection (Walsh and Lynch 2018), but the actual values are not crucial for the main argument of this paper. This results in a Gaussian fitness curve, with the variance in the number of offspring being 0.90, resulting in an opportunity of selection (variance in relative fitness) being *I* = 0.14, meaning that the phenotype can at most change with √*I* = 0.38 standard deviations (Walsh and Lynch 2018). The fitness of the individual is thus the fitness propensity of each individual. W_max_ was set such that the population size remained approximately stable over time, in this setting 1.7.

Individuals were then paired by splitting the population into two halves, where the first half was mated to the second half, i.e. random mating with regard to phenotype. The number of offspring of each pair was determined by the sum of the fitness values of each individual in each pair, rounded to nearest integer, as offspring tend to come in wholes rather than fractions. Each offspring was given a genotype by randomly assigning one allele from each parent at each loci. Hence, there is a variance among offspring due to recombination, as in any sexually reproducing species. All parents die after mating and the offspring constitutes the new population. There was no inbreeding in any of the populations (mean F = -0.0046, SD = 0.0071, range: -0.026, + 0.013).

In total 100 populations with the same start as above were exposed to fluctuating selection for 100 generations. The fluctuations in the optimum were random with no temporal autocorrelation within a range of ± 1 SD of the mean. Each population was independent from all other populations in terms of environmental fluctuations. After the 100 generations the phenotypic means were significantly different (P < 0.001, effect size (eta-squared) = 15.5 %, min = 143.7 (se = 0.17), max = 155.4 (0.22)). There was no correlation between original mean phenotype and the subsequent response to the selection below (r^2^ = 0.2 %).

After 100 generations an experiment was made where for each population the optimum changes 1.0 standard deviation from the mean. This is a strong directional selection corresponding to a variance standardized selection gradient on average 0.37. The response in phenotypic mean was predicted in three different ways. First, the classic Breeder’s equation R = h^2^S was used, where S = cov(*z,w*), where *w* is relative fitness calculated as the number of offspring divided by the mean number of offspring. The selection gradient is then 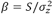(Walsh and Lynch 2019). I also used the traditional Lande-Arnold linear regression incorporating stabilising selection: w = a + β_1_z + β_2_z^2^ + ε, using mean-centered phenotypic values (z). However, the results did not differ from the Breeder’s equation, indicating no influence from stabilizing selection, and these values are therefore not shown.

Second, I used the correction for unequal reproductive success (Walsh & Lynch 2019, pp 487-488),

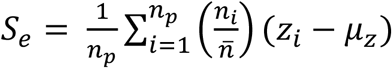

Where n_p_ is the number parents breeding, n_i_ is the number of offspring of the ith pair, and z_i_ is the mean phenotypic value of the parents, and 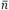is the mean number of offspring. The selection gradient using this approach will be called the Effective selection gradient, β_Eff_.

Third, I used the mean fitness and mean phenotype of the parents. This was done to cover the fact that it is the pair that is breeding unit and not single individuals in a sexually reproducing species, and hence the offspring are a function of the combination of males and females. With this approach the additive genetic variance needs to be adjusted to be the variance in mean parent breeding values. In both cases I used the mean-centered male and female values (z_i_ – zmean). The estimation of predicted was then carried out in the same way as above, but with the mean values instead of individual values. The selection gradient using this approach will be called the Pair selection gradient, β_Pair_

The path to the new optimum (θ) can be approximated by a logistic function of mean value (Fig 2) as a function of time (t):

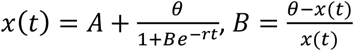

The population is assumed to be at optimum at the inflection point of the function, which is the time when the curve becomes flat, and can be found as -Log(B)/r, where *r* is estimated from the data All analyses and simulations were made in Mathematica 12.0.0.0 (Wolfram Research Inc. 2019).

## Results

On average, the experiment resulted in a change in the predicted direction. However, there was a considerable variation around the mean, with around 10 % in the wrong direction for the 100 independent populations (Fig 1). There was a large within-population variation, i.e. the same population differ in the response if we replicate the experiment exactly (Fig 1, Single), and the same was true if h^2^ = 1 (Fig 1, Genotypes). If we run the 100 population 100 times each then the variance among populations in the observed response was found to be 9.6 %. If we set the heritability 1.0 (i.e. phenotypes equal the genotypes), we still find a large variance among populations (70.8 %), but now the within-population variance is relatively larger (29.2 %), as compared to the case when the heritability was 0.5 (9.6 %).

**Figure 1.**
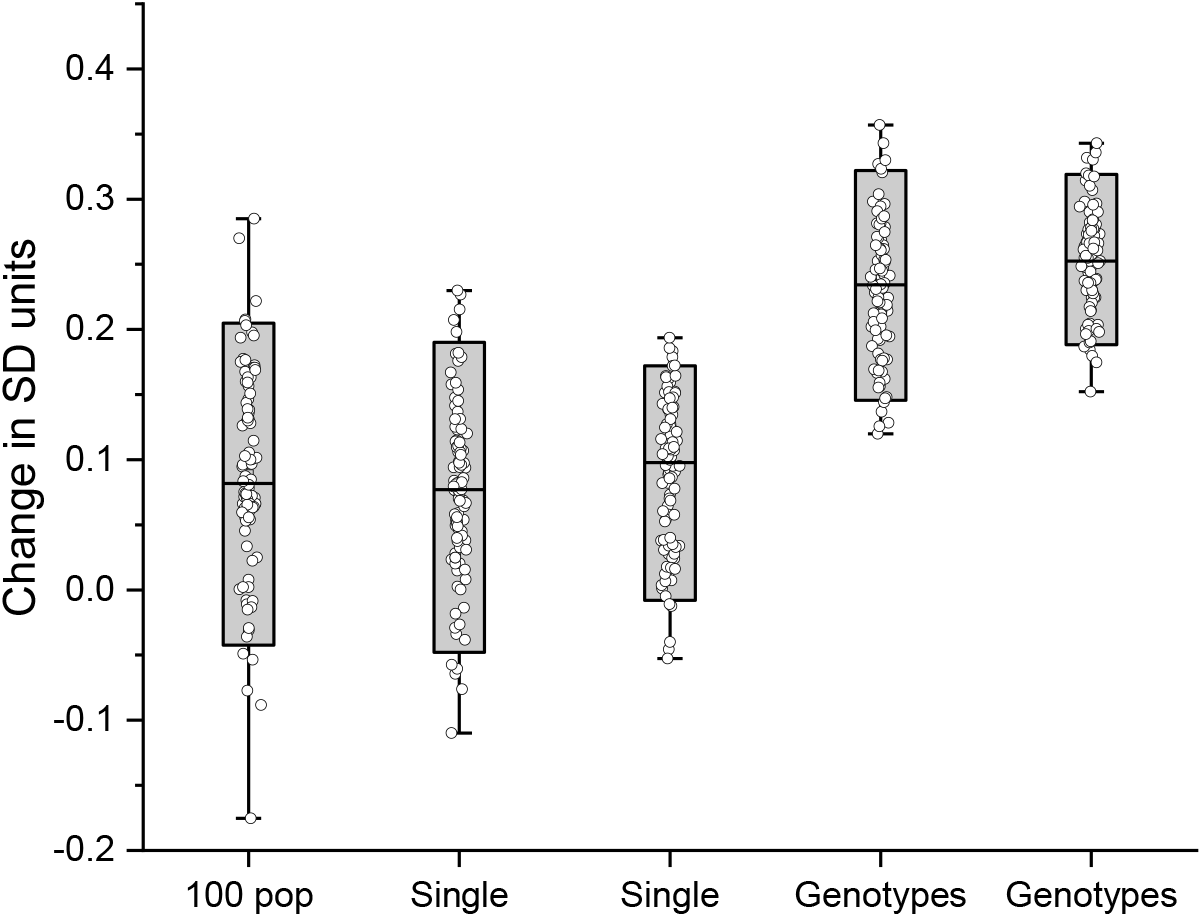
The observed response for 100 separate populations, for two populations replicated 100 times each (Single), and for two populations replicated 100 times each with a heritability of 100 % (Genotypes).

The variation among runs can also be seen in the time to reach the new optimum (Fig 2a). If we use 100 different populations, the time ranges between 10 and 140 generations (median 43-44 generations, SD = 19.4 and 18.5, for the two runs respectively, Fig 2b). The probability of reaching the new optimum at 25-30 generations is as high as between 55-60 generations (15%). There is a variance in trajectories and in inflection time even when the heritability is set to 100 % and for the same population with the same selection (Fig 2c and d).

**Figure 2.**
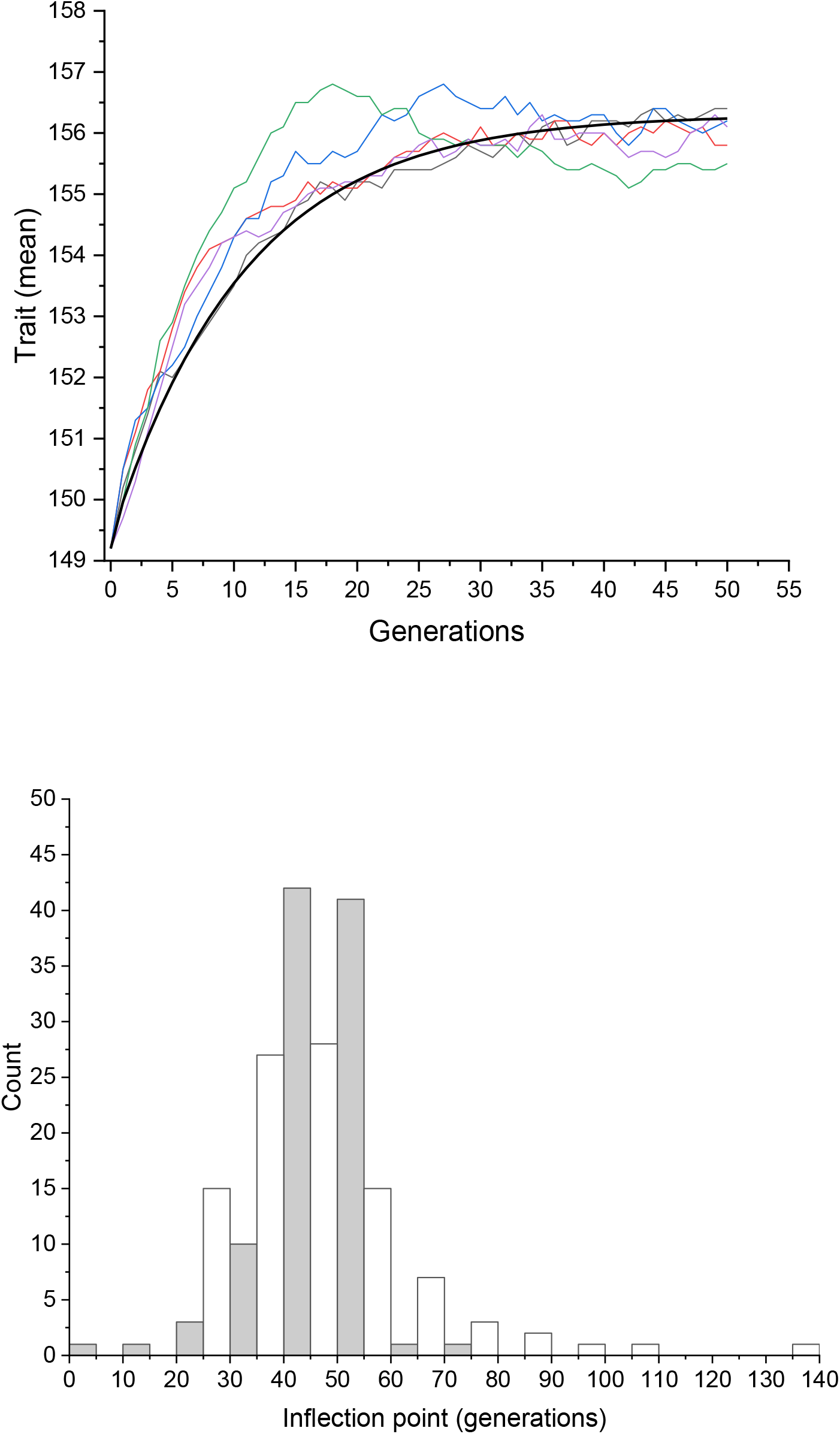
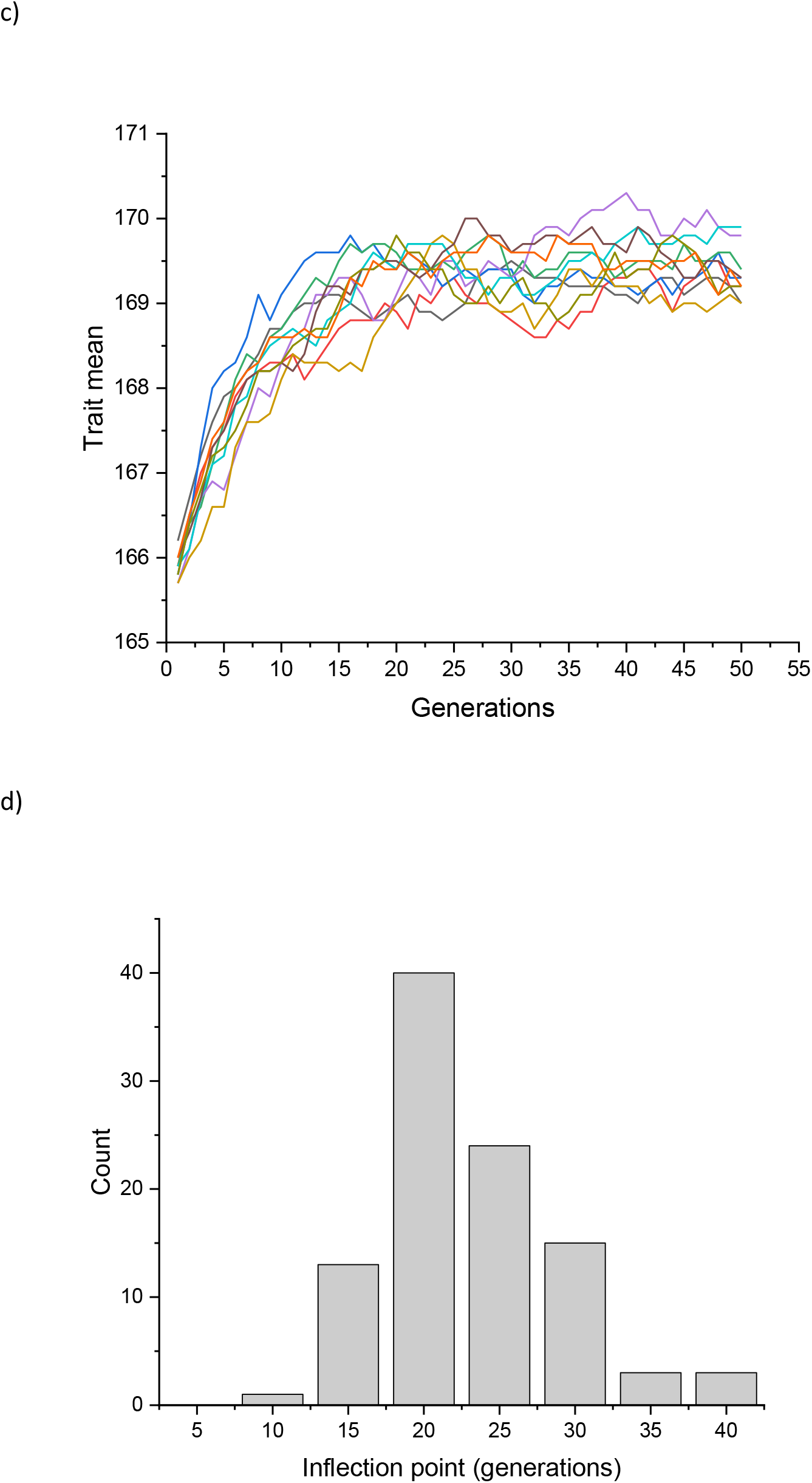
Time to reach the optimum. A) five examples, with a theoretical curve fitted from the data. B) distribution of inflection points from two sets of 100 populations, and c) trajectories using one population with 100 % heritability d) distribution of inflections points for the populations with 100 % heritability

Second, the different methods to predict change differ greatly in their accuracy, with the Breeder’s equation consistently overestimating the response, and greatly so, and the Pair-approach is still biased upwards but not greatly so. The bias in response can be seen in Fig 3 using all data where 95 % of the predicted responses using the Breeder’s equation were biased. For the Effective selection approach 71 % of the predicted responses were larger than observed, while for the Pair-approach 56 % of the predicted response was larger than observed. There was also a difference in precision (measured as 1/σ^2^) with the Pair-approach being seven times more precise than the Breeder’s equation (95.6 vs. 13.8), and three times more precise than the Effective selection (95.6 vs 31.9). The Effective selection was in turn 2.3 times more precise than Breeder’s equation.

**Figure 3.**
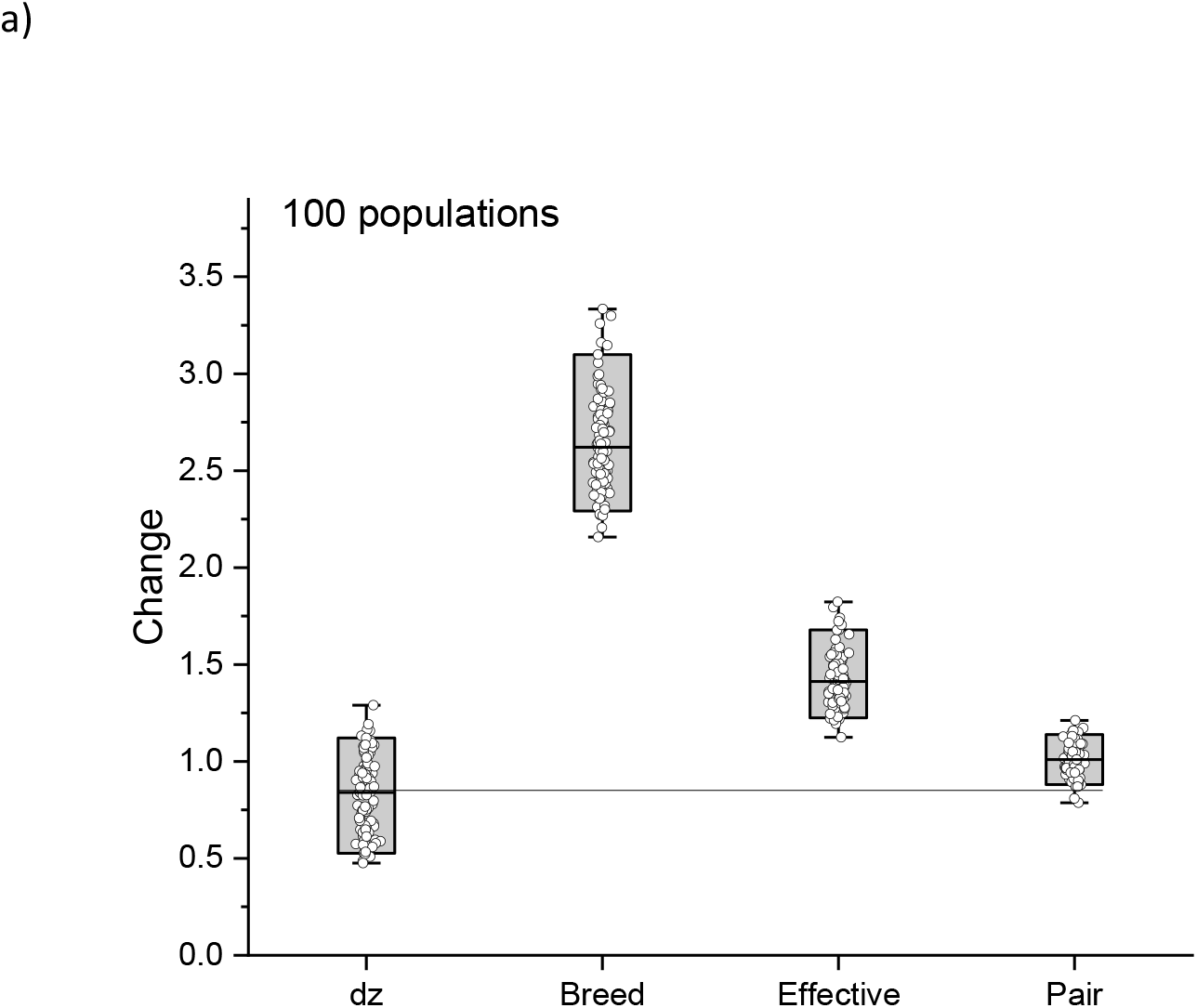
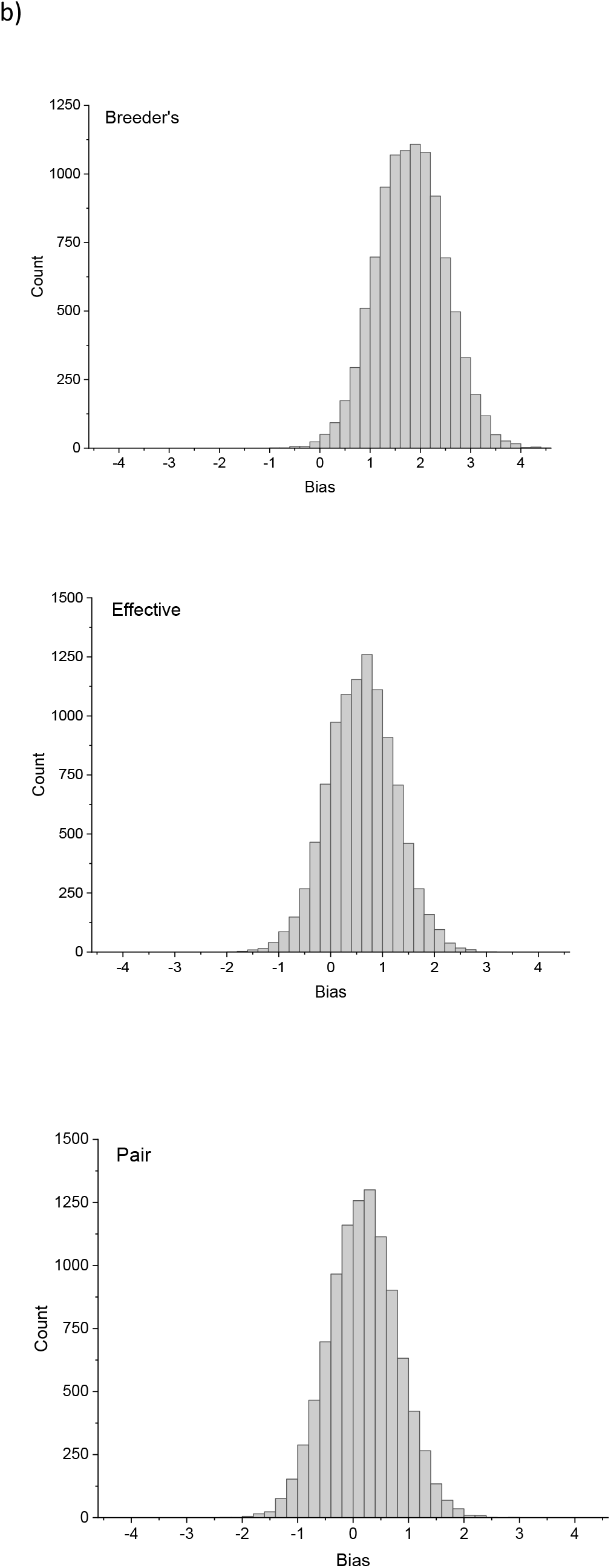
a) The observed and predicted change using three different methods. b) distribution of bias (predicted – observed) using the three different methods to predict the response.

To get an understanding of the source of the bias we can look more closely to the two parts of the predicted response, selection gradient and additive genetic variance. The selection gradient was largest for β_Pair_, followed by β and with β_Eff_ as the smallest. In all populations was β_Pair_ larger than β, which means that the difference in selection gradients cannot explain the difference in predicted response (Fig 4a). The difference between β and β_Pair_ was significant in all populations (P << 0.001, Wilcoxon test). However, there was a large difference in genetic variance, where V_a_ using the individual approach was about three times larger than V_A_ measured at the pair level (mean (range) over populations Va = 29.42, range 25.65-34.84, V_APair_ = 10.60, range 8.62-13.04, Fig 4b).

**Fig 4.**
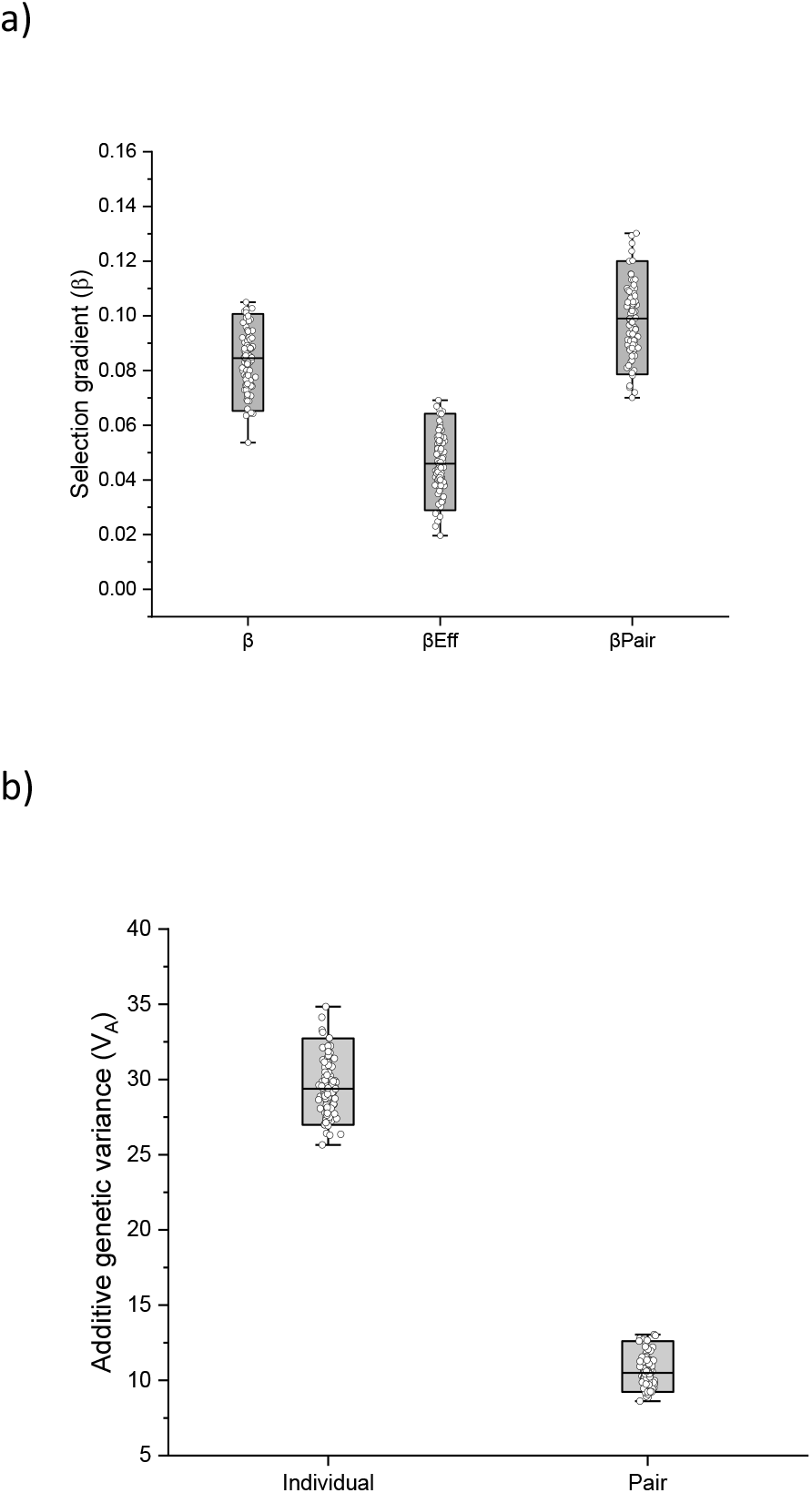
a) selection gradients for the three methods. b) Additive genetic variance using the variance of individuals breeding values (Individual), and the variance in the mean pair breeding values (Pair).

## Discussion

There are two major results from this study. First, there is a large variance of outcomes for each selection event, even with identical initial conditions and with identical origin. This variation is due to the different history of the populations (variance among populations), the imperfect genotype- phenotype map (phenotypic variance within populations) and the variance in offspring genotypes due to recombination and segregation (variance among genotypes within populations). Note that the whole population of 500 individuals was used and hence the variance is not due to sampling. Second, the estimation of genetic variance needs to be made on the level of the pair since not doing so can lead to a significant upward bias of the of the predicted response. All these factors are likely to contribute significantly to the mismatch between the predicted and observed response (e.g. Pujol et al. 2018)

The results show a great variation among replicate runs using exactly the same population and the same selection even if we bypass the phenotypic level. This show that the response to selection is a unique event that cannot be replicated exactly. In a diploid population with sexual reproduction the number of genotypes in the offspring is related to the number of loci and increases very rapidly to astronomically large numbers. This means that at any given reproductive event with a finite number of offspring the probability of getting two offspring with the same genotype is essentially zero. Combined over a number of pairs we can see that a variance in outcome will happen by necessity. It is a fundamental consequence of sexual reproduction. This is nothing new as it is well-known that the expected genotype of an offspring of pair of individuals is the mean of the genotypic values with a variance equal to half the variance in genotypic values between the parents (Bulmer 1980). It just seems to be forgotten in the discussion of the response to selection.

The variance in outcome was apparent at the genotypic level, and magnified to the phenotypic level, which is a necessary consequence of the simple genotype-phenotype map used here. Exactly how this map looks like will differ between populations and is an empirical question that is related to the environment where the population lives. The many studies of phenotypic plasticity have shown that the connection between genotypes and phenotypes is not obvious and nonlinearity is not uncommon (e.g. West-Eberhard 2003). This will add to the lack of match between predicted and observed outcomes, especially if the environment changes faster and in the opposite direction of selection (Merilä et al. 2001b). This adds to the difficulties in predicting the phenotypic response. Note that hard to predict does not mean impossible; there are cases in the literature where a good match between predicted and observed response has been found even if they are not as common as we would like it to be (Walsh and Lynch 2018; Pelabon et al. 2021).

Finally, the variance among populations means that the history of selection for each population matters for the response to a new selection. The role of history (historicity), has been stressed repeatedly for many years (e.g. Lewontin 1966, Beatty and Dejardins 2009; Desjardins 2011). This poses a number of empirical challenges since for the vast majority of species and populations basically nothing is known about their selective past. Yet it can play a role for the future response. At the time being the only thing we can do is to keep this in mind when predicting future evolutionary change.

All these three factors work together to create a variance around the expected response to selection. Thus, the expected evolutionary response should be seen as the mean (expectation in statistical terms) of a distribution of values with a variance determined by the underlying genetic variances and covariances, the genotype-phenotype map and the history of selection. The fact that there is an uncertainty of in the estimate of the predicted response due to sampling is known (Pelabon et al. 2021), while, for example, the variance due to recombination and its stochastic segregation of alleles into the offspring needs far more attention. This means that even if the additive genetic variance, V_A_, is identical for two populations the response will most likely differ between the populations even if the selection is the same. In fact, the same population will not respond in the same way twice. It is important to remember that V_A_ is a summary statistic that has important evolutionary implications (e.g. Hansen and Pelabon 2021), but for the prediction of a response to selection from one generation to the next in a given situation it is simply not precise enough. The variance of outcomes needs to be considered.

The predicted response to selection using the variance in individual breeding values resulted in an upward bias, while using the combined pair breeding values resulted in almost unbiased predictions with the highest precision. This is an example of the regression to the mean originally exposed by Galton (1886). An extreme individual will not have extreme offspring since in every sexually reproducing species it has to mate with someone that is probably less extreme. This means that the offspring will be a compromise between the two genotypes, a classic result of quantitative genetics (Bulmer 1980). This suggests that the unit of selection and V_A_, when it comes to predicting future response, is the reproducing pair (see Björklund and Gustafsson 2013). This can be estimated with least problems in monogamous systems where pair stay together, but can get far more complicated to estimate in other mating systems, for example when there is multiple paternity. This is an empirical issue that should not be underestimated. The variance in the mean breeding values of the pair was found to give the most accurate prediction of the response, while using a more individual-based approach grossly overestimate the genetic variance that matters for the evolutionary change.

The results in this study has an implication that is very important for the following discussion. Two populations with a common ancestor will diverge genetically, *by necessity* (e.g. Brandon and McShea 2020). This will in turn have an effect on their future evolutionary trajectories. If the evolutionary response is determined by unique population and time specifics, then each response will be unique. Two responses might very well be very similar, and not different in a statistical test, but they will never be identical. It comes down the details of the genetics of the traits under study, i.e. the number of alleles segregating, which in turn is a result of past selection and drift, and the structure of the genotype-phenotype map. The vast combinations of possible combinations of genotypes has a very important implication; each population is a unique collection of genotypes, a collection that has not been seen before, and will never be seen again. This means that every reproductive event gives rise to a new collection of genotypes being different from every other reproductive event by a very high probability. This uniqueness of sexually reproducing populations suggests an open-ended evolution, and hence posing substantial problems in terms of predicting the future as suggested by Day (2013).

The results have implications for empirical studies where a set of populations are compared in relation to some ecological factor. A consensus has emerged that one needs a number of spatial replicate population to test if factor X leads to response Y. Theoretically this is correct and mimics what is done in the lab. However, this is not as simple in nature. An assumption of replicates is that they should not differ more than trivially, something that is clearly violated in nature since populations will be divergent by their unique history of selection and recombination. In addition, given the capricious outcome of a reproductive event due to recombination and segregation the number of replicate populations needed will by necessity be very large. In short, most studies will be underpowered and often greatly so. As the results show, populations selected to reach a new optimum will reach the optimum in due time, but the variance in the time to reach the new optimum can differ substantially. This in turn means that if we get inconsistent results from a study of natural population we simply need much more data and more time to see the pattern, if there is one. A recent example of two closely related species with the same selective environment but with divergent response can, for example, be found in Hartke et al. (2020).

The results have a bearing on the discussion on the “replication crisis” in science. Given that populations differ and the capricious nature of recombination should we really expect populations respond in the same way given the same selection? The fact that certain results cannot be replicated in nature might not be the problem, the problem is instead the assumption that we can expect results to be replicated in different natural populations (barring foul play, of course). Turning things on this side suggests that we should *not* expect results to be replicated other than in special cases. Differences might be the rule, rather than similarities. So rather than expecting similar response and being worried about lack of it, we should expect different responses and be thrilled (or worried) whenever consistent results are found. The idea that experiments should be replicated in natural populations comes from the lab world where this can be expected given the huge possibilities to standardise groups. However, this might apply only to the same lab populations as different labs can still get different results even if standardisation has been taken to a maximum (e.g. Ackerman et al. 2001).

## Conclusions

The evolutionary process is messy and is rarely straightforward. This is a fundamental feature as a result from the stochastic nature of history, selection, recombination and segregation. This study shows that this can have a profound effect on our ability to predict evolutionary change. The predicted response should thus be seen in terms of a distribution of responses with a mean and variance. The study also shows that the traditional method of estimating genetic variance using individual breeding values leads to an overestimation of the predicted response. On the other hand, using the variance of the mean breeding values of each pair leads to an essentially unbiased prediction.

## Acknowledgements

None

## Literature cited

Beatty, J. and E. C. Desjardins. 2009. Natural selection and history. Biol. Philos. 24: 231–246.

Björklund, M., and L. Gustafsson. 2013. The importance of selection at the level of the pair over 25 years in a natural population of birds. Ecol. Evol. 3: 4610–4619.

Brandon, R.N., and D. W. McShea. 2020. The missing two-thirds of evolutionary theory. Cambridge Elements, The Philosophy of Biology. Cambridge Univ Press. Cambridge, UK.

Bulmer, MG. 1980. The mathematical theory of quantitative genetics. Clarendon Press, Oxford, UK.

Chatfield, C. 2004. The analysis of time series. 6th Ed. Chapman & Hall/CRC. London, UK.

Day, T. 2013. Computability, Gödel’s incompleteness theorem, and an inherent limit on the predictability of evolution. J. R. Soc. Interface 9: 1–16.

Desjardins, E. 2011. Historicity and experimental evolution. Biol. Philos. 26: 339–364.

Dingemanse, NJ., Araya-Ajoy, Y., and D. Westneat. 2021. Most published selection gradients are underestimated. Why this is so and how to fix it. Evolution 75: 806–818.

Galton, F. 1886. Regression towards mediocrity in hereditary stature. J. Anth. Inst. Great Britain and Ireland. 15: 256–263.

Hansen, T.F., and C. Pelabon. 2021. Evolvability: a quantitative-genetics perspective. Ann. Rev. Ecol. Evol. Syst. 52: 153–175.

Hartke, J., Waldfogel, A-M., Sprenger, P.P., Schmitt, T., Menzel, F., Pfenniger, M., and B. Feldmeyer. 2020. Little parallelism in genomic signatures of local adaptation in two sympatric, cryptic sister species. J. Evol. Biol. 34: 937–952.

Heywood, J.S. 2005. An exact form of the Breeder’s equation for the evolution of a quantitative trait under natural selection. Evolution 59: 2287–2298.

Lynch, M., and B. Walsh. 1998. Genetics and analysis of quantitative traits. Sianuer, Sunderland. MA. USA.

Merilä, J., Sheldon, B. C., and L. E. B. Kruuk. 2001a. Explaining stasis: microevolutionary studies in natural populations. Genetica 112: 199–222.

Merilä, J., Kruuk, L. E. B., and B. C. Sheldon. 2001b. Natural selection on the genetic component of variance in body condition in a wild bird population. J. Evol. Biol. 14: 918–929.

Merilä, J., and A. P. Hendry. 2014. Climate change, adaptation and phenotypic plasticity: the problem of evidence. Evol. Appl. 7: 1–14.

Morrissey, M.B., Parker, D.J., Korsten, P., Pemberton, J.M., Kruuk, L.E.B., and A. J. Wilson. 2012. The prediction of adaptive evolution: empirical application of the secondary theorem of selection and comparison to the Breeder’s equation. Evolution 66: 2399–2410.

Pelabon, C., Albertsen, E., Le Rouzic, A., Firmat, C., Bolstad, G., Armbruster, W.S., and T. F. Hansen. 2021. Quantitative assessment of observed versus predicted response to selection. Evolution 75: 2217–2236.

Pujol, B., Blanchet, S., Charmantier, A., Danchin, E., Facon, B., Marrot, P., Roux, F., Scotti, I., Teplitsky, C., Thomson, C. E., and I. Winney. 2018. The missing response to selection in the wild. Trends Ecol. Evol. 3: 337–346.

Walsh, B., and M. Lynch 2018. Evolution and selection of quantitative traits. Oxford University Press. Oxford, UK.

West-Eberhard, M.J. 2003. Developmental Plasticity and Evolution. Oxford University Press: New York, USA.

Wolfram Research, Inc. 2019. Mathematica, Version 12.0.0.0, Champaign, IL.

